# Measuring Growth, Resistance, and Recovery after Artemisinin Treatment of Plasmodium falciparum in a single semi-high-throughput Assay

**DOI:** 10.1101/2024.11.11.623064

**Authors:** Mackenzie A.C. Sievert, Puspendra P. Singh, Douglas A. Shoue, Lisa A. Checkley, Katelyn M. Brenneman, Tarrick Qahash, Zione Cassady, Sudhir Kumar, Xue Li, François H. Nosten, Timothy J.C. Anderson, Ashley M. Vaughan, Jeanne Romero-Severson, Michael T. Ferdig

## Abstract

**Background:** Artemisinin partial resistance (ART-R) has spread throughout Southeast Asia and mutations in *pfKelch13*, the molecular marker of resistance, are widely reported in East Africa. Effective *in vitro* assays and robust phenotypes are crucial for monitoring populations for the emergence and spread of resistance. The recently developed extended Recovery Ring-stage Survival Assay used a qPCR-based readout to reduce the labor intensiveness for *in vitro* phenotyping of ART-R and improved correlation with the clinical phenotype of ART-R. Here, we extend and refine this assay to include measurements of parasite growth and recovery after drug exposure. Clinical isolates and progeny from two genetic crosses were used to optimize and validate the reliability of a straight-from-blood, SYBR Green-based qPCR protocol in a 96-well plate format to accurately measure phenotypes for Growth, Resistance, and Recovery.

**Results:** The assay determined growth between 6 h and 96 h, resistance at 120 h, and recovery from 120 h and 192 h. Growth can be accurately captured by qPCR and is shown by reproduction of previous growth phenotypes from HB3 × Dd2. Resistance measured at 120 h continually shows the most consistent phenotype for ring stage susceptibility. Recovery identifies an additional response to drug than parasites that are determined sensitive by Fold Change at 120 h. Comparison of progeny phenotypes for Growth vs Resistance showed a minor but significant correlation, whereas Growth vs Recovery and Resistance vs Recovery showed no significant correlation. Additionally, dried blood spot (DBS) samples matched Fold Change measured from liquid samples demonstrating Resistance can be easily quantified using either storage method.

**Conclusions:** The qPCR-based methodology provides the throughput needed to quickly measure large numbers of parasites for multiple relevant phenotypes. Growth can reveal fitness defects and illuminate relationships between proliferation rates and drug response. Recovery serves as a complementary phenotype to resistance that quantifies the ability of sensitive parasites to tolerate drug exposure. All three phenotypes offer a comprehensive assessment of parasite-drug interaction each with independent genetic determinants of main effect and overlapping secondary effects that should be further. By adapting our method to include DBS, readouts can be easily extended to *ex vivo* surveillance applications.

## Background

Malaria is endemic to most tropical regions with Sub-Saharan Africa bearing the largest burden accounting for 94% of the roughly 249 million cases and 608,000 deaths worldwide (1). Current front-line drug treatments for uncomplicated *falciparum* malaria rely on Artemisinin (ART) derivatives due to their effectiveness on circulating ring-stage parasites and rapid reduction of patient parasitemia (2). Artemisinin partial resistance (ART-R) spread throughout the Greater Mekong Subregion of Southeast Asia, largely mediated by mutations in the Kelch propellor domain of the gene *pfKelch13* (K13) (3). Many resistance-conferring mutations have arisen independently and repeatedly in the region making K13 status a reliable marker for resistance surveillance in Southeast Asia (4). The recent emergence of ART-R parasites in multiple East African countries with and without K13 mutations raises concerns about the prolonged efficacy of ART on the African continent (5–7). Consequently, adequate surveillance for the emergence and spread of ART-R in Africa will require more than knowledge of K13 mutations. Comprehensive measurements of a wider profile of ART-R phenotypes will support more effective surveillance.

Clinical and *in vitro* measurements of ART-R are both standardized to important components of assessing rapid clearance the pharmacological hallmark of ART (8). The clinical phenotype for ART-R, Parasite Clearance half-life (PC_1/2_), is the most direct measurement and is defined as the time needed for parasite load to be reduced by half, and it is measured as part of highly coordinated therapeutic efficacy studies (9). This approach is instrumental for detecting reduced drug efficacy and the emergence of resistance, but is logistically difficult and expensive, limiting its use (10,11). *Ex vivo* or *in vitro* assays directly assess and confirm the parasite susceptibility to drug by removing patient factors such as immunity, spleen activity and drug absorption that can confound the *in vivo* phenotype of the parasite (9,12,13). In resource limited areas, *ex vivo* or *in vitro* assays are a cost-effective way to generate baseline drug susceptibility data to direct efficient implementation of therapeutic efficacy studies. Clinical and *in vitro* phenotypes work in tandem to inform national malaria control programs, emphasizing the need for an improved throughput, information rich *in vitro* assay that can accurately capture quantitative levels of drug resistance and is accessible to resource limited or remote sites for routine surveillance (10).

The Ring-stage Survival Assay (RSA) has been widely adopted as the most effective ART-R *in vitro* correlate of PC_1/2_ (14). The RSA mimics the pharmacokinetics of ART by exposing a highly synchronous culture of early ring-stage parasites to a clinically relevant concentration of dihydroartemisinin (DHA) for 6 h. The RSA phenotype is reported as percent survival, calculated by dividing the parasitemia of a drug treated culture by the parasitemia of an untreated culture 66 h following drug removal. The RSA is technical and labor intensive, requiring large culture volumes, parasite synchronization, and determination of viable vs non-viable parasites by either visual inspection of blood smears under a light microscope or highly specialized flow cytometry protocols. The microscopy-based readout increases accessibility of the assay and offers more flexibility for processing experiments but introduces reader subjectivity into the phenotype, impacting results and interpretation. Whereas flow cytometry-based readouts are less subjective but must be completed immediately at the conclusion of the 66 h incubation in RSA and require specialized equipment and expertise on site. Technical difficulties associated with RSA limit throughput of the assay thereby reducing the amount of information gained by *ex vivo* surveillance or *in vitro* experiments.

Previous studies using Southeast Asian parasites occasionally demonstrate discordant phenotypes between PC_1/2_ and RSA with no clear explanation for the discrepancy between the *in vitro* and *in vivo* phenotypes (14,15). Identification of parasites with discordant phenotypes have also been observed in a much smaller sampling of African patient samples, although patient immunity is overall greater in African which could influence the relationship between these phenotypes (16). These inconsistencies highlight an opportunity to expand the *in vitro* assay to capture a broader range of the parasite response to drug. Early investigations to define ART-R examined a variety of approaches to identify a strong *in vitro* correlate with PC_1/2_: 72 h growth inhibition, 48 h growth inhibition, 24 h trophozoite maturation inhibition, trophozoite-stage survival assay, mature-stage survival assay, recovery assay, ring-stage growth arrest assay, quiescence survival assay, and a delayed clearance assay to evaluate ART-induced dormancy (8,17–24). These various phenotypes captured an array of potential mechanisms that could influence clinical ART-R by broadly assessing physiological characteristics of drug response observed by ART-R parasites. In particular, assays for quiescence and dormancy examined an aspect of ART-R that have been hypothesized to contribute to clinical treatment failures and been recently show to be present in human infections following artesunate therapy (25,26). However, these phenotypes were not easily quantified into a single measurement and carried challenges to optimization needed for surveillance-scale throughput. None of these assays incorporated an aspect of parasite fitness, a key component of evolutionarily successful resistance with potential to spread. A deeper analysis of parasite phenotypes of ART efficacy may provide better predictors of emergence of resistance and even may provide an explanation for distinct responses among different parasites. This will be especially beneficial in areas of emerging resistance where there is no guarantee ART-R will evolve identically to Southeast Asia.

Previously, we used 15 clinical isolates with known PC_1/2_ phenotypes to develop and optimize the extended Recovery Ring-stage Survival Assay (eRRSA) to increase the throughput and accuracy of the *in vitro* measurement for ART-R (27). eRRSA is reported as Fold Change, as the assay employed a straight-from-blood qPCR protocol to measure the fold-change difference in DNA content between DHA-treated and untreated cultures. To best match PC_1/2_, incubation time after drug removal was also extended to 114 h, compared to 66 h in RSA, allowing an additional cycle of active proliferation for all cultures. ART-R parasites in DHA-treated cultures proliferate comparably to the untreated cultures and thus, produce lower fold-change differences than sensitive lines. The requirement for active growth of treated cultures to produce a lower Fold Change ensures a measurement of viable parasites which more accurately measures the true level of ART-R. This refined assay and quantitative readout generates a large separation between sensitive and resistant lines as well as stronger correlation with the clinical phenotype than the traditional RSA percent survival readout (27). The streamlined qPCR methodology and Fold Change readout increases throughput by eliminating the need for a laborious microscopy or expensive and specialized flow cytometry-based readouts. Samples can also be frozen for long-term storage which offers much more flexibility for later quantification. The ease of sample collection and processing afforded by the qPCR platform allows for expansion of the assay to analyze more phenotypes through the collection of samples at additional timepoints.

In the current study we expand on our eRRSA approach to include measurements for innate parasite Growth rate as well as parasite Recovery after drug treatment. The new phenotypic measurements incorporate qPCR values into a single plate-based, semi-high throughput Growth, Resistance, and Recovery Assay (GRRA). Using the Growth, Resistance, and Recovery Assay (GRRA) we show that parasite growth and recovery are quantitative traits that segregate (i.e. are inherited differently) across progeny of the HB3 × Dd2 and NF54(HT-GFP-luc) × NHP4026 genetic crosses. Additionally, we show the ART-R phenotype is readily determined from dried blood spots (DBSs) of DHA-treated and untreated cultures in side-by-side comparison with qPCR from matched liquid samples. Using DBS samples for long term preservation and sample accumulation is an advantage for low-resource clinical or field sites by eliminating the requirement for freezers and qPCR equipment to be on-site. The adaptability, flexibility, methodological ease, relatively low cost, and series of quantitate phenotypes measured in the GRRA will provide a path to new insights into the functional mechanisms responsible for differential parasite responses to ART exposure.

## Methods

### Parasite lines

Cloned parasite lines for this study encompass parental parasites of recent experimental genetic crosses (KH004-H9, MAL31, MKK2835, NHP1337, NF54HT-GFP-luc, and NHP4026) (28), an additional ART sensitive isolate (NHP4302), two multidrug resistant lines (VN-C1 and VN-E10), as well as select progeny from the NF54(HT-GFP-luc) × NHP4026 cross and select progeny of the HB3 × Dd2 cross. Clinical isolates were chosen based on known ART-R phenotypes and used to validate the optimal sampling timepoints. The progeny from NF54(HT-GFP-luc) × NHP4026 were chosen for further examination due to the Resistance and Recovery phenotypes of the parents and to quantitatively assess the relationship between the Growth, Resistance, and Recovery phenotypes (29,30). Parents and progeny from HB3 × Dd2 were chosen for their known growth rates as a means to validate the growth measurement component of the GRRA (31).

### Thawing of Parasite Stocks

Cryovials containing the parasites frozen in glycerolyte 57 (Baxter Healthcare) were removed from the freezer or liquid nitrogen, thawed at room temperature, and the contents were transferred to a 50 mL centrifuge tube. An equal volume of 12% NaCl, was slowly added dropwise to the tube with agitation between drops. A solution of 1.67% NaCl was then slowly added dropwise to the 50 mL tube to a volume of 5 mL and then centrifuged at 680x*g* for 5 m and the supernatant removed. The final thawing solution, 0.9% NaCl and 0.2% dextrose, was slowly added dropwise to the tube to a volume of 5 mL and centrifuged again at 680x*g* for 5 m. The supernatant was removed and the RBC pellet containing both infected and noninfected cells was suspended in 3 mL of complete media at 5% hematocrit then transferred to a 25 cm^3^ flask.

### Culturing conditions

*Plasmodium falciparum* parasite lines were cultured using standard laboratory protocols and maintained at a Hematocrit of 5%. Flow cytometry was performed on a Guava easyCyte (described below) to estimate parasitemia and to adjust cultures as needed to maintain parasitemia ≤ 3%.Thin blood smears were made daily to monitor the health of parasites. Red blood cells (RBCs) (Biochemed Services, Winchester, VA and Interstate Blood Bank, Memphis, TN) were washed 3X using equal volumes of incomplete media (ICM)(RPMI 1640 with L-glutamine (Gibco, Life Technologies.), 50 mg/L hypoxanthine (Calbiochem, Sigma-Aldrich), 25 mM HEPES (Corning, VWR)). Washed RBCs were suspended as a 50/50 mixture in ICM prior to being used in cultures. Complete media used in cultures was made by adding 0.5% Albumax II (Gibco, Life Technologies.), 10 mg/L gentamicin (Gibco, Life Technologies) and 0.225% NaHCO_3_ to incomplete media. Parasites were grown in flasks at 37°C under a 5% CO_2_/5% O_2_/90% N_2_ gas mixture.

### Flow cytometry

To determine stage and parasitemia, 80 μL was removed from each culture, centrifuged at 376x*g* for 1 m, and supernatant removed. Pellets were washed with 200 μL of flow wash buffer (FWB) consisting of a 10% Fetal Bovine Serum (FBS) solution in RPMI without phenol red (Thermo Fisher). Samples were centrifuged again at 376x*g* for 1 m, the supernatant was discarded, and the pellet was suspended in 80 μL of FWB. 25 μL of sample was mixed with 25 μL of dye master mix (2μL of a 5X SYBR + 2 μL of 62.5 μM Syto61 in 21 μL of FWB) and incubated at 37℃ for 40 m. Samples were then washed 3x with 200 μL of FWB and pellets suspended in 100 μL of FWB. Eight μL of stained sample was added to 200 μL of FWB in individual wells of a 96-well plate and the 96-well plate was loaded on to a Guava easyCyte HT. 50,000 events were counted to estimate parasitemia, and percent ring stage calculated based on gated cells. As a control to assess the staining quality and ensure accurate gating, an uninfected red blood cell culture was maintained identically to parasite cultures and was included in all steps of the protocol with the parasite samples.

### Growth, Resistance and Recovery Assay protocol

Parasite cultures that were predominantly segmented schizonts were synchronized using a single percoll layer method (32). Briefly, 8 mL of cultures were centrifuged at 900x*g* for 4 m and supernatant removed. The remaining RBC pellet was suspended in 2 mL of incomplete media and then gently layered on to a 3 mL aliquot of percoll (Sigma Aldrich) diluted with 1X RPMI (Thermo Fisher) and 13.3% sorbitol (Sigma Aldrich) in phosphate buffered solution (Fisher Scientific) to a concentration of 70% in 15 mL centrifuge tubes. Layers were then centrifuged at 1575×*g* for 10 m with no brake. The top layer containing RBCs with late stage schizonts was moved into a new 15 mL tube and rinsed 3x with incomplete media by centrifuging at 900x*g* for 4 m. Supernatant was removed between rinses. The resulting pellet was resuspended in 2 mL of complete media at 2% hematocrit and placed on a shaker at 37℃ for 4 h. At the 4 h mark, 80 μL was removed from the culture and used for flow cytometry determination of stage and parasitemia as described above. Cultures that met quality control requirements of ≥ 70% early ring stage and ≥ 1% parasitemia were adjusted to 0.5% parasitemia at 2% hematocrit. Treated samples were dosed with 700 nM DHA in 0.02% Dimethyl sulfoxide (DMSO); untreated samples were exposed to an equivalent amount of 0.02% DMSO. Master mixes of samples were aliquoted into 3 technical replicates of 275 μL each in 96 well plates (3 treated, 3 untreated). At 6 h post-exposure, 225 μL of media was removed from the wells and replaced with 225 μL of incomplete media and spun for 2 m at 800x*g*. The Rinse was repeated a total of 3 times after which 225 μL complete media added back to the wells. Thin microscopy smears were made using 2 μL of culture for each sample and 20 μL of culture was transferred to PCR strips and stored at -80°C until qPCR quantification. Additionally, 20 μL of culture samples and thin microscopy smears were collected at 96, 120, and 192 h post-treatment. At 120 h, media was changed on all wells and fresh complete media added back.

### Determination of optimal sampling times for multiple phenotypes

Nine recent clinical isolates (NHP4026, NHP4302, NHP4373, NHP1337, MKK2835, KH004-H9, MAL31, VN-E10, VN-C1) and NF54HT-GFP-luc were evaluated in a 10-day eRRSA time series experiment to establish the minimal number of timepoints required to gain the growth, resistance, and recovery phenotypic information in one assay. Parasites were synchronized, drug treated, and culture growth was monitored for 10 d. 20 μL of culture was frozen for qPCR quantification and thin smears were made for microscopic evaluation of treated and untreated cultures daily for the first 144 h and then every second day until 240 h.

### qPCR quantification and phenotype calculations

A standard curve of known parasitemia was produced using the NF54 parasite line for each qPCR quantification for conversion from Cycle threshold (Ct) to parasitemia and comparison between qPCR experiments. The culture was grown to 10 mL then synchronized and at 18 h post-synchronization, stage and parasitemia were evaluated by flow cytometry. The culture was adjusted to 5% parasitemia at 2% hematocrit and used to generate an 8-step, 3-fold dilution series. Each step of the standard curve was separated into 100 μL aliquots in PCR strips and frozen at -80 °C. At the time of qPCR experiments, aliquots of the standard curve and 20 μL culture samples were removed from -80 °C, allowed to thaw at room temperature and mixed well. Samples and the standard curve were amplified by qPCR using the Phusion Blood Direct PCR kit (ThermoFisher, cat # F547L). Each individual reaction mix contained 3 μL of sample as DNA template, 0.1 μL of Phusion enzyme, 5 μL of 2X Phusion Blood Direct Mix Buffer, supplemented with 1.4 μL of 7.5X SYBR Green in DNase/RNase free water (ThermoFisher) and 0.25 μL each of forward and reverse primers from a 10 μM stock (designed to amplify the highly conserved region of *pfCRT*) for a final volume of 10 μL per qPCR (Additional file 1). Cycling conditions were as follows: 20 s denaturation at 95 °C, followed by 30 cycles of 95 °C for 1 s, 62.3 °C for 30 s, and 65 °C for 15 s on an ABI 7900HT thermocycler (Additional file 1). Ct was calculated using ABI SDS 2.4.1 software. For Fold Change (2^ΔCt^), ΔCt is calculated using the average Ct of the treated replicate samples minus the average Ct of the untreated replicate samples in the following equation:

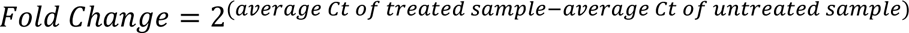

Parasite Multiplication Rate (PMR) to quantify parasite growth was then assessed from the untreated samples taken at 6 h and 96 h (33). Ct values were converted to parasitemia using the line equation generated from the standard curve included in each qPCR experiment. PMR was calculated using the average parasitemia for untreated samples at 6 h and 96 h post-treatment in the following equation:

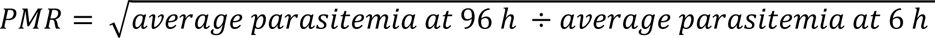

Finally, Recovery following drug treatment was quantified using samples from DHA-treated samples at 120 h and 192 h post-treatment. Ct values were converted to parasitemia using the standard curve, averaged, then used in the following equation to calculate Recovery:

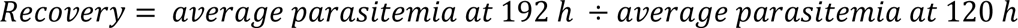

### Sampling dried blood spots from filter papers

Seven clinical isolates and six clonal unique progeny lines with known resistance phenotypes were measured for Fold Change at 120 h using liquid samples and DNA eluted from samples spotted on filter paper. DBS samples were made by spotting 8 μL of sample onto Whatman grade 3 qualitative filter paper (VWR, cat #1003-917) cut to 96-well plate size. Liquid culture samples were pipetted carefully drop-wise to keep the circumference of the spot as small as possible, roughly 7mm large, to facilitate uniform and complete excision of the DBS in a single punch. Filter paper samples were dried and stored in a sealed plastic bag at ambient temperature until processing. The 7mm diameter filter paper samples were placed in 0.1 mL PCR strips and 50 μL of Phusion blood direct 2X buffer added to fully submerge to the DBS in buffer. The PCR strips were moved to a shaking incubator platform at 60℃ and set to 180 rpm for 30 m. Samples were removed and visually inspected for complete removal of the DBS from the filter paper. PCR strips were centrifuged at 2000×*g* for 1 m and the supernatant removed to fresh PCR strips. PCR strips containing filter paper punches were centrifuged a second time and additional eluted buffer was removed to the new PCR strip to maximize sample recovery. The supernatant was slightly discolored due to elution of the DBS. The sample supernatant was used for qPCR reactions as described previously, with slight modifications to reaction procedure as follows: 5 μL of sample supernatant was added to a 5 μL of qPCR mix containing 0.25 μL each of both forward and reverse primers from 10 μM stock, 0.1 μL Phusion enzyme, 1.4 μL 7.5X SYBR Green in DNase/RNase free water, and an additional 3 μL of pure DNase/RNase free water. The number of cycles was increased to 40 while all other cycling conditions remained the same.

### Data Analysis

Briefly, progeny and parents of the NF54(HT-GFP-luc) × NHP4026 genetic cross were assayed in three blocks with each block containing parental genotypes; the use of replicated parental genotypes allowed for detection of block effects. Analysis of variance was used as described previously to identify technical variation in Growth, Resistance, and Recovery phenotypes amongst progeny and parents(30,34). To account for the technical variation confounding the examination of biological variation, all phenotype values within each block were divided by the mean phenotype value of the control parent, NF54HT-GFP-luc. This procedure effectively normalized the average NF54HT-GFP-luc phenotype to 1 while maintaining an estimation of error for the parental phenotype and allowed for comparison of progeny phenotypes relative to the control parent. All further statistical tests and linear regressions were performed in the graphing and statistics software GraphPad Prism 10 (Version 10.1.2).

## Results

### Optimal sampling times for multiple phenotypes

Fold Change was tracked over 10-days which confirmed 120 h as the optimal timepoint for *in vitro* assessment of ART-R. Evidenced by consistent measurements of sensitive parasites and best distinction of resistant parasites from the qPCR readout (Fig. 1A, Additional file 2)(27). Fold Change values at 120 h for all nine parasites distinguished sensitive, resistance and moderately resistant parasites (Additional file 3). The time series also demonstrated that ART sensitive parasites Fold Changes decrease after 120 h, with NHP4026 showing the sharpest decrease by 192 h (Fig. 1A, Additional file 2). To more closely examine this drop in Fold Change after 120 h, DHA-treated and untreated samples were separated for NF54HT-GFP-luc and NHP4026 to assess how the DNA content of each culture influenced Fold Change (Fig. 1B). The untreated samples of both parasites plateau after 144 h, indicating that cultures have grown to capacity; however, DNA content is not decreasing and therefore not driving the drop in Fold Change (Fig. 1A). The DHA-treated samples of both parasites increased in DNA content, demonstrating that the culture has begun to recover from treatment and proliferate by 192 h post-treatment (Fig. 1B). The increase in DNA content of the DHA treated culture for both parasites explains the drop in Fold Change at 192 h observed in Fig. 1A. Recovery was assessed for ART sensitive clinical isolates by comparing 192 h relative to 120 h for three isolates from Southeast Asia, NHP4032, MKK2835, and NHP4026 and two African isolates, NF54 and MAL31. The Southeast Asian isolate NFP4026 recovered significantly more quickly after drug exposure compared to either African isolate (Fig. 1C).

**Fig. 1.**
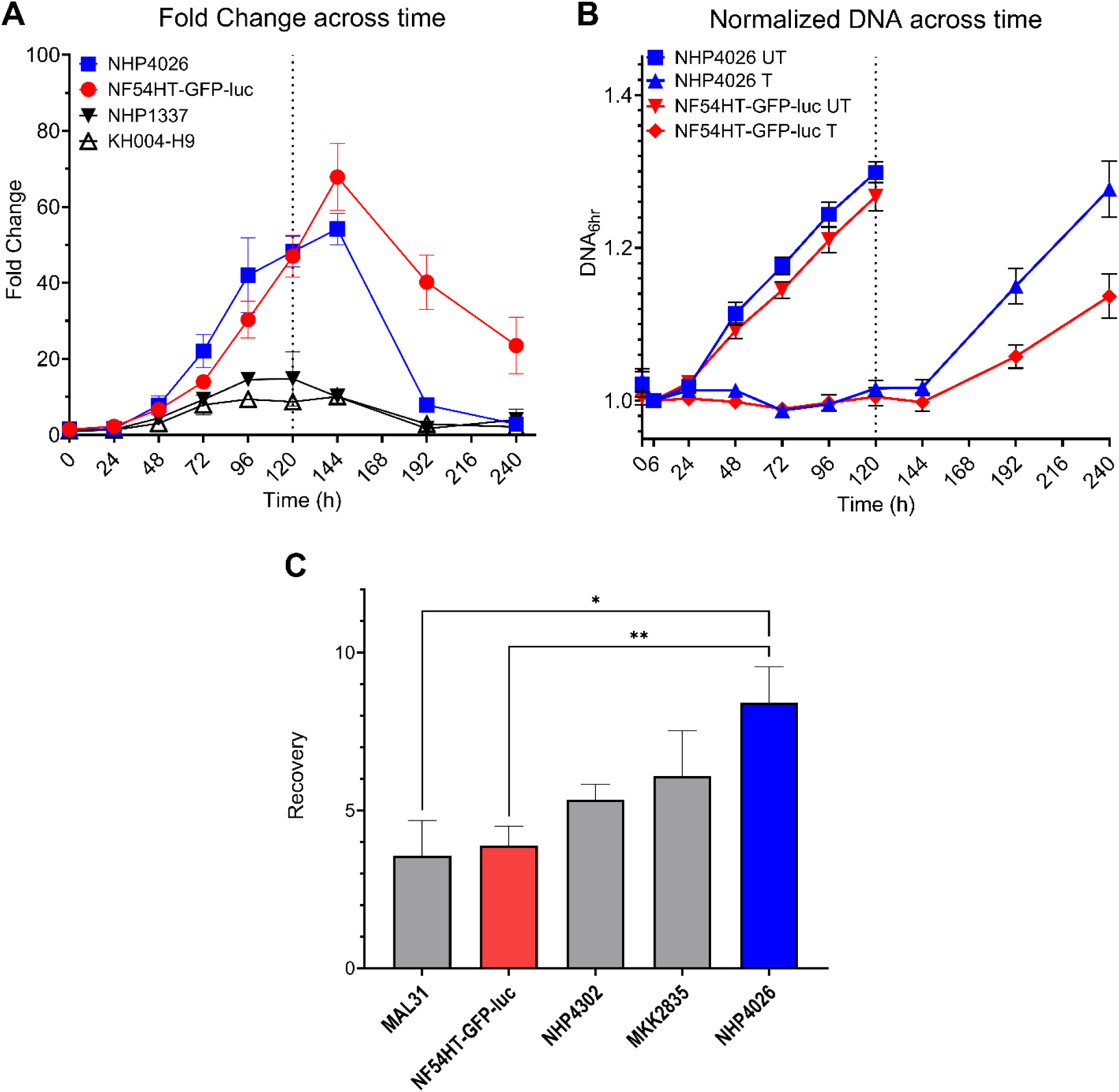
Fold Change and DNA content of clonal *Plasmodium falciparum* lines across time. **A)** Fold Change in DNA content between DHA treated and untreated samples of representative artemisinin resistant and sensitive parasites across the 10-day time series. **B)** DNA content of DHA treated (T) and untreated (UT) cultures NHP4026 (Blue) and NF54HT-GFP-luc (Red) across the 10-day time series is shown normalized to DNA content at the 6 h timepoint. **C)** Quantification of Recovery after drug treatment measured in four recent clinical isolates and NF54HT-GFP-luc. Data is reported as mean ± SEM of at least three biological replicates. Statistical significance between parasites was determined by one-way ANOVA (\**P* < 0.05; \*\**P* < 0.01).

### Quantification of the Recovery phenotype in artemisinin sensitive parasites

We further explored the potential for Recovery to be a reliable, quantitative phenotype revealing a novel aspect of the drug-parasite interaction distinct from RSA or eRRSA. We measured this trait in 23 ART sensitive progeny derived from the experimental cross between NF54(HT-GFP-luc), an African ART susceptible strain and NHP4026, a Southeast Asia isolate that demonstrated a strong slow clearing clinical phenotype but ambiguous *in vitro* resistance (29,35). Establishing the inheritance of this trait among progeny clones derived from these parents would confirm that genetic differences between them influences their recovery phenotype (Fig. 2). The progeny show a quantitative distribution with NHP4026 at the high end of the distribution, as would be expected if this is genetically determined; only 2 progeny had more rapid recovery (Fig. 2). NF54HT-GFP-luc is centered within the distribution and more than half of the progeny exhibited lower mean Recovery (Fig. 2). Importantly, the Resistance and Recovery phenotypes of the parents and ART sensitive progeny are not significantly associated (R^2^ = 0.0021, F(1, 23) = 0.0501, *P*-value = 0.8249) (Additional file 4), suggesting that the main genetic drivers of the two phenotypes are not correlated and indicating these traits are determined by distinct genetically inherited mechanisms.

**Fig. 2.**
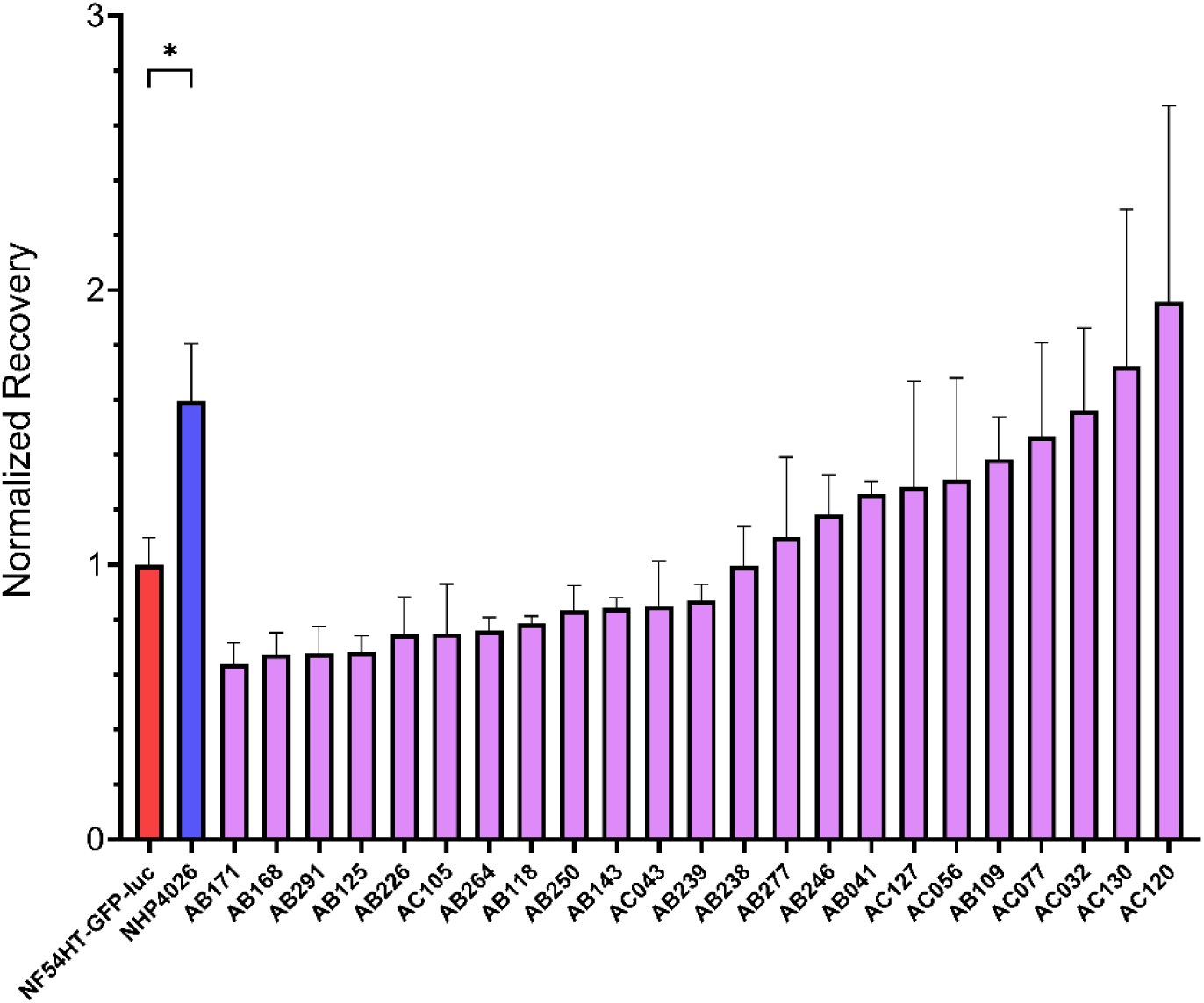
Distribution of Recovery phenotypes for ART sensitive parasite lines. Recovery after drug treatment measured in 23 progeny from the NF54(HT-GFP-luc) × NHP4026 genetic cross with phenotype values normalized to the NF54HT-GFP-luc parent. Data is reported as mean ± SEM of at least three biological replicates. Statistical significance between the parents NF54HT-GFP-luc and NHP4026 was determined by unpaired Student’s *t*-test (\**P* < 0.05).

### Parasite Multiplication Rate (PMR) estimated from qPCR can distinguish growth rate differences between parasites

We used a subset of eight progeny and both parents from HB3 × Dd2 to validate PMR derived from our qPCR readout (Fig. 3A). As expected, Dd2 proliferates at a significantly higher rate than HB3 (unpaired Student’s t-test, p-value = 0.03) and supported observations from our earlier studies measuring growth related phenotypes that included increase in parasitemia across eight days, asexual cell cycle duration, merozoites per schizont, and red blood cell invasion efficiency (36). We further used simple linear regression to examine the relationship between PMR and asexual (erythrocytic) cell cycle duration in progeny of HB3 × Dd2 (31). The overall regression for the two phenotypes was statistically significant (R^2^ = 0.4661, F(1, 8) = 6.984, *P*-value = 0.0296), which illustrated that parasites with shorter cell cycles proliferate faster (Additional Figure 5). However, the progeny SC01 deviated from the regression line more than any other parasite due to a lower PMR phenotype than what would be expected given the cell cycle duration. Regardless, quantifying PMR using qPCR reflects the trends for known innate growth rate differences of these parasite lines and can serve as a reliable approximation of parasite growth as an additional phenotype derived from our assay.

**Fig. 3.**
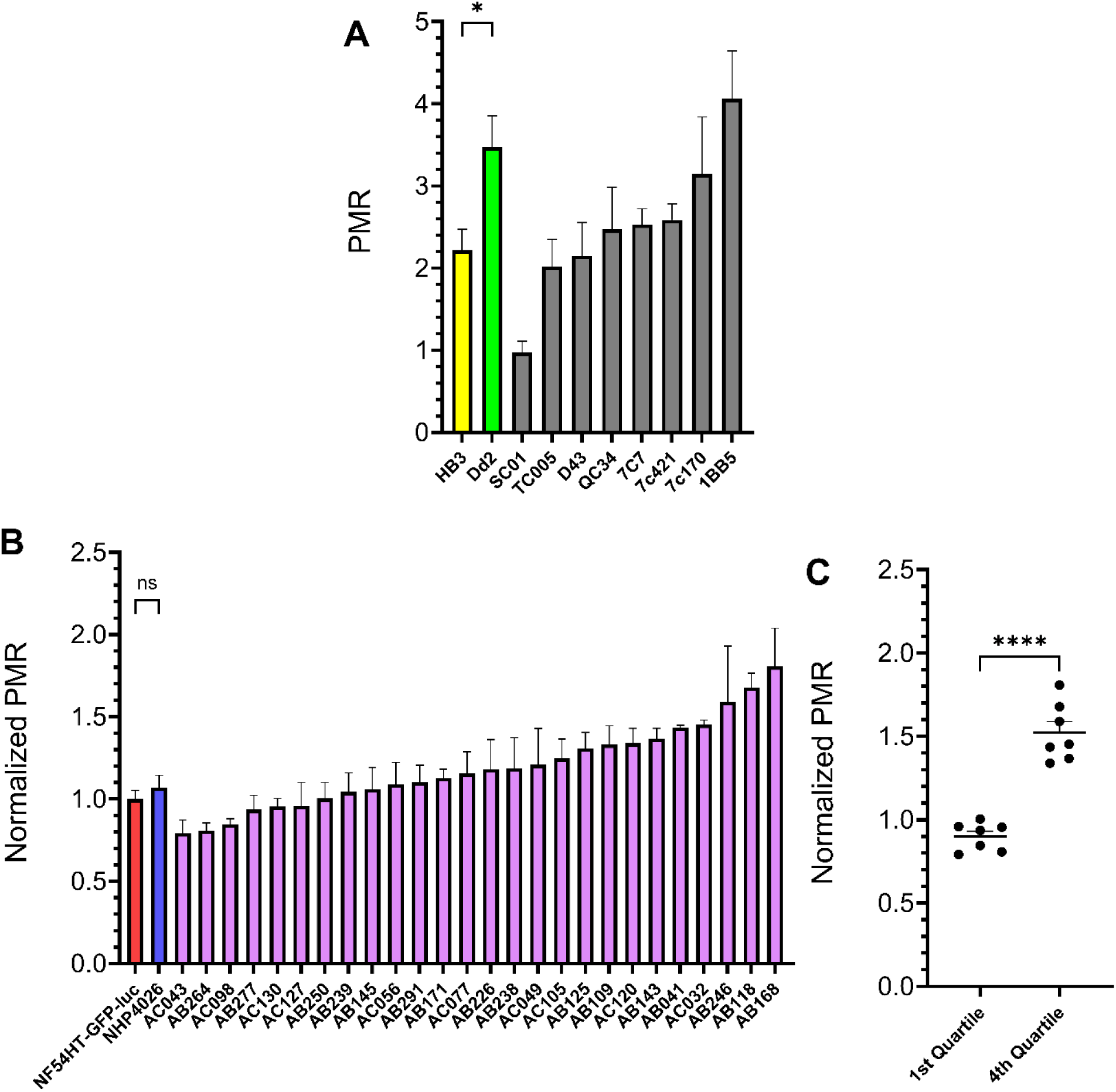
Parasite Multiplication Rate (PMR) of progeny from two experimental genetic crosses. **A)** PMR for eight progeny and parents of the HB3 × Dd2 genetic cross. **B)** PMR for 26 progeny and parents of the NF54(HT-GFP-luc) × NHP4026 genetic cross with phenotype values normalized to the NF54HT-GFP-luc parent. **C)** Comparison between phenotype means of progeny in the first and fourth quartile of the NF54(HT-GFP-luc) × NHP4026 genetic cross phenotype distribution shown in **B**. Data is reported as mean ± SEM of at least three biological replicates. Statistical significance was determined by unpaired Student’s *t*-test (\**P* < 0.05; \*\**P* < 0.01; \*\*\**P* < 0.001; \*\*\*\**P* < 0.0001; ns, not significant).

We also examined PMR in progeny and parents of NF54(HT-GFP-luc) × NHP4026 to assess whether parasite growth rates may affect drug response readouts. The parent lines, NF54HT-GFP-luc and NHP4026, have nearly identical growth rates and the majority of progeny have a mean PMR only slightly greater than the parents (Fig. 3B). However, comparison of the progeny with the most extreme PMR phenotypes in the first and fourth quartile show a significant difference between the ends of the phenotype distribution (Fig. 3C). Isolation of progeny with extreme phenotypes suggests that a combination of alleles at multiple loci inherited from both parents can increase PMR beyond the parental phenotypes. Further, we observed a significant positive linear relationship between Fold Change and PMR (R^2^ = 0.2317, F(1, 27) = 8.141, *P*-value = 0.0082) indicating that 23% of variation is shared between the phenotypes (Additional file 6). The co-linear relationship between Resistance and Growth is consistent with parasites with higher levels of resistance are less fit and grow slower compared to highly sensitive parasites. Notably, there is no significant relationship between Recovery and PMR (R^2^ = 0.001, F(1, 27) = 0.0289, *P*-value = 0.8662), indicating that innate parasite growth rate does not predict that rate of recovery after drug exposure (Additional file 6).

### Resistance phenotypes assessed using dried blood spots on filter paper

Resistance phenotypes measured in parallel from the same parasite isolates and progeny based on liquid or filter paper were highly correlated (Spearman r = 0.87, *P*-value = 0.0002, Fig. 4). The progeny AC105 is the only parasite that exhibited a large deviation in Resistance between the two methods (Fig. 4). However, by both methods AC105 is highly ART sensitive relative to the other parasites measured. Both liquid and filter paper stored samples accurately identified highly resistant, moderately resistant, and sensitive parasites with consistent interpretation.

**Fig. 4.**
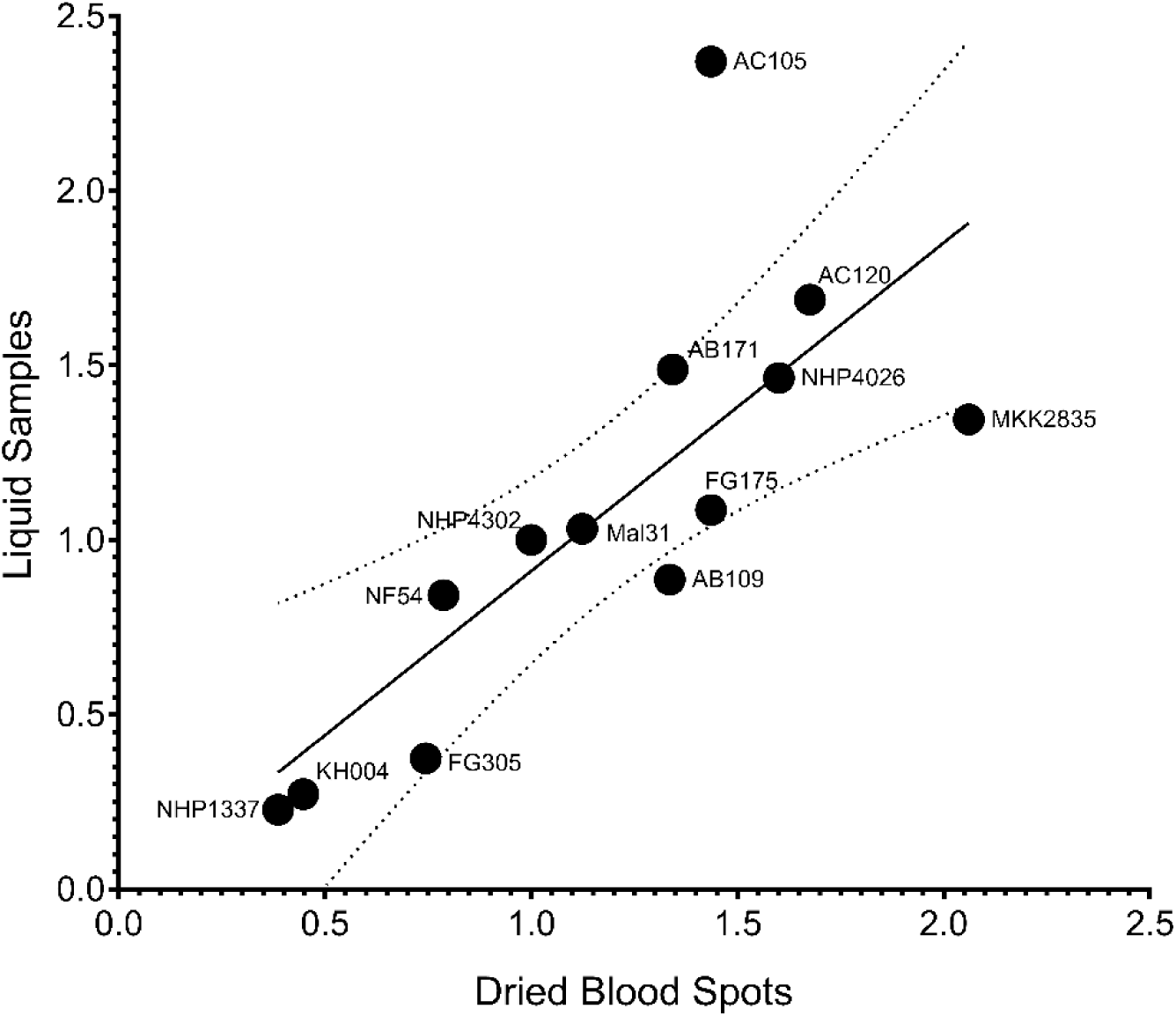
Scatterplot and correlation of Resistance phenotypes for 13 parasite lines generated from dried blood spots vs liquid samples. At 120 h post-drug treatment samples from whole culture were spotted onto filter paper as dried blood spots or frozen at -80°C as liquid samples. Fold Change values measured from DNA eluted from DBS were correlated with Fold Change values measured from matched liquid samples (Spearman r = 0.8721, *P*-value = 0.0002) and the 95% confidence interval around the best-fit line (*y* = 0.94*x* – 0.02927) is denoted with dotted lines. Data was normalized to the ART sensitive control parasite NHP4302 and each parasite phenotype is represented as the mean of at least 3 biological replicates.

## Discussion

Here we report and validate a single, easily accessible assay to quantify multiple phenotypes that are relevant to the increasing challenge of surveilling ART-resistance emergence and spread. Current phenotypic surveillance of ART resistance relies on the laborious and time-consuming RSA, an assay requiring tight synchronization, time sensitive drug dosing, and expert examination of blood smears using light microscopy to assess live vs dead parasites. Here we have expanded on our earlier development of eRRSA (27) as a streamlined and quantitative alternative to traditional RSA, to now assay novel phenotypes including rate of recovery after DHA exposure as well as an innate growth rate phenotype. The speed and ease of the qPCR readout relative to microscopy or flow cytometry enables both the throughput and standardization. It also allows for the modification of time points, doses, and other variables to generate new insights into parasites’ responses to drug with minimal additional labor and expense, not amenable to traditional microscopy.

By conducting an eRRSA time series experiment we were able to confirm 120 h post-DHA exposure is the optimal time point at which to quantify ART resistance using a qPCR readout, originally benchmarked against PC_1/2_ (27). Furthermore, we observed that some ART sensitive parasites as determined by standard eRRSA recover more rapidly after drug treatment. Therefore, we sought to quantify enhanced Recovery as a reproducible phenotype that could be used to detect early or emerging low-level resistance, perhaps as an initial step in the evolution of high-level resistance.

The three Southeast Asia ART sensitive isolates in our initial testing panel exhibited an enhanced Recovery phenotype compared to NF54HT-GFP-luc and the African isolate, Mal31 (Fig. 1C). We observed significant differences in Recovery between the parents NF54HT-GFP-luc and NHP4026 for an initial suggestion of a genetic basis for the phenotype (Fig. 1C). We leveraged a subset of progeny from NF54(HT-GFP-luc) × NHP4026 to examine the pattern of inheritance underlying Recovery. NHP4026 is one of the most rapidly recovering parasites with only two progeny with a greater Recovery whereas NF54HT-GFP-luc is centered in the phenotype distribution with 13 progeny phenotyped with a lower mean Recovery (Fig. 2). A phenotype distribution with progeny showing phenotypes more extreme than the parents is suggestive of transgressive segregation, where alleles from both parents contribute genetic determinants that influence the Recovery phenotype through complex genetic interaction. The progeny used in this proof-of-concept experiment demonstrate that Recovery is a reproducible and quantitative phenotype suitable for association studies and linkage mapping. However, due to the relatively small number of progeny, this screen lacks the statistical power for an unbiased, genome-wide linkage analysis to identify the interactions between loci. The progeny set for this cross has been expanded by additional selection and cloning to now encompass 270 unique recombinant progeny (30) and we eagerly anticipate conducting a large mapping study using GRRA to identify loci underlying Recovery (37).

Interestingly, NHP4026 does not have a Resistant *in vitro* phenotype by Fold Change (Fig. 1, Additional file 2 and 3), but does have a slow clearance clinical phenotype (8.37 PC_1/2_) (38) and a previously reported resistant RSA (≥ 10% survival) (35). The Resistance and Recovery data provides an opportunity to investigate different features of drug response in this atypical ART-R parasite. NHP4026 has the highest Recovery out of all isolates tested and is nearly at the top of the distribution of Recovery phenotypes for NF54(HT-GFP-luc) × NHP4026. We suggest that a quiescence or dormancy phenotype could explain the discordant *in vitro* and *in vivo* phenotypes for NHP4026. Quantification of RSA and PC_1/2_ rely heavily on assessment of the morphology of the drug-treated parasites, and NHP4026 parasites treated with DHA could present morphology of viable ring stage parasites but not be actively proliferating. However, to produce a resistant measured by Fold Change, parasites in treated samples must be actively proliferating in the additional life cycle up to the 120 h timepoint (Additional file 2, Additional file 3). Therefore, GRRA, producing both Fold Change and Recovery results in the same assay could give more refined information to assess “intermediate” parasites that are not strongly ART-R or ART-S but may be critical for monitoring resistance emergence. Interestingly, Resistance and Recovery are not significantly correlated in this progeny set, indicating that a major genetic determinant(s) is not shared between these phenotypes (Additional file 4), and we anticipate that a large-scale screen of the NF54(HT-GFP-luc) × NHP4026 could provide insights into the mechanisms controlling both phenotype and evolutionary path to ART-R.

Temporary growth arrest after treatment with clinically relevant doses of ART derivatives, described as quiescence or ART-induced dormancy, has been hypothesized to contribute to clinical treatment failures of ART monotherapy resulting in recrudescent infections. Assays have been developed to specifically examine quiescence and ART-induced dormancy, but require additional experimental steps post-treatment to remove non-dormant parasites or to track parasite growth by fluorescent imaging (24,39). These additional steps preclude the throughput required to screen large numbers of parasites and cannot be easily integrated into the GRRA. Recovery as measured by GRRA is distinct from a true dormancy assay in that there is not a second purification of the culture after treatment to remove any parasites that do not enter a dormant state. Instead, the GRRA identifies parasites that recover rapidly after DHA treatment. Parasites with higher Recovery will define a small pool for deeper examination of quiescent or dormant phenotypes.

Drug resistant parasites often acquire compensatory mutations to counteract fitness costs of resistance mutations. Well compensated parasites present a greater risk of spreading through a population. Parasite growth rate has been used as a proxy for fitness and is an important initial measure to characterize parasites in areas of emerging resistance. We demonstrated in 8 progeny and parents from the HB3 × Dd2 that PMR can be effectively measured using qPCR and can recapitulate known growth differences between parents and progeny (31,33). Linear regression of PMR with previously measured asexual cell cycle duration shows the two phenotypes are moderately associated, with 46% of variation in PMR explained by asexual cell cycle duration (Additional file 5). Nine of 10 parasites fit this relationship well with one, SC01, deviating from the trend line. The low number of total parasites used in the comparison may overestimate the relatedness of the two phenotypes, however SC01 could be an outlier for PMR due to other asexual growth phenotypes shown to be significantly different between HB3 and Dd2 (36). We anticipate that mechanisms underlying PMR are diverse and specific to an individual parasite or an individual population of parasites. PMR as part of GRRA offers an initial screen to identify resistant, fast-growing parasites that may indicate higher relative fitness to be explored in directed subsequent experiments.

Examination of PMR in NF54(HT-GFP-luc) × NHP4026 provided insight into the possible relationships between growth and drug response phenotypes in the genetic background of a recent clinical isolate. Parents of this cross have nearly identical PMRs which could initially suggest the underlying genetic determinants influencing PMR are identical in the parents and would ultimately be an uninformative phenotype to measure in progeny of this cross (Fig 3B). However, using only a subset of progeny we identified a range of PMRs including a significant difference between parasites with the highest and lowest PMR (Fig. 3C). This indicates that progeny at the extreme ends of the phenotype distribution inherited alleles from different loci contributing to higher PMR in each parent which combined as synergistic epistasis in some progeny. The increase in PMR is significant but relatively small and would likely require many progeny to detect the interaction at a statistically significant threshold.

Regression of PMR and Fold Change showed a significant linear association with 23% of variation shared between the phenotypes (Additional file 6). It is generally thought that resistant parasites proliferate more slowly due to the cost of resistance mutations that impact biological processes required for growth. This could explain the significant linear relationship; however, we note that the relatively low predictive power of growth by resistance level suggests that the overlap of determinants controlling each phenotype is partial at best. This is not surprising because many different pathways can impact growth rates regardless of their association with resistance.

Importantly, the linear relationship between PMR and Recovery was not significant, suggesting the genetic control of these two phenotypes are independent of each other (Additional file 6). It might be assumed that parasite growth rate would dictate its expansion rate after drug exposure, however, our progeny phenotypes do not support that. Rather, it seems that Recovery as assessing parasite growth only between 120 h to 192 h after drug exposure emphasizes how long it takes parasites to resume normal growth.

Linkage studies investigating a phenotype in separate genetic crosses is a powerful approach to dissect the underlying genetic architecture of genes influencing a phenotype of interest. The potential of this approach is highlighted by PMR measured here in HB3 × Dd2 and NF54(HT-GFP-luc) × NHP4026. HB3 and Dd2 are near the extremes of the phenotype distribution, indicative of a progeny set suited to identify loci with large effects independent of non-additive gene interactions. Conversely, NF54HT-GFP-luc and NHP4026 are centered within the progeny distribution with nearly all progeny with higher or lower PMR, indicative of a progeny set suited to identify a complex architecture of multiple and interacting loci with smaller effects. The success of genetic cross experiments rely on prior knowledge of the parent phenotypes to develop informative crossing schemes to test specific hypotheses. The incorporation of PMR into routine surveillance will develop this knowledge about clinical isolates to identify strong candidate parasite isolates for further *in vitro* experiments or as parents of future experimental genetic crosses.

While widespread use of GRRA in clinical settings is constrained by the requirement of parasite culturing facilities and freezers, our ability to spot liquid culture samples onto filter paper for long term preservation and sample accumulation will allow for more efficient qPCR. Identical samples measured from liquid and DBS from filter paper correlated strongly (Spearman r = 0.87, *P*-value = 0.0002) and obtained consistent interpretation of Resistance for every parasite tested (Fig. 4). Our observations are consistent with recent reports using a dry-preservation method to store samples from *ex vivo* IC_50_s of multiple drugs (40). Importantly the two approaches diverge in important ways with the *ex vivo* IC_50_ quantification relying on PfHRP2 ELISA which touts quantification of very low-level parasitemia using inexpensive plate readers. Nevertheless, reagents can be expensive and ART-R parasites with *PfHRP2* deletions are increasing in prevalence in African countries. Comparatively, the qPCR-based method would only require adjustment of primers rather than developing new antibodies, a much less expensive and faster assessment of the level of any single-copy gene. Moreover, filter paper samples can be easily shipped to any site with an adequate qPCR machine for processing and data generation, eliminating the need for equipment on-site. Both methods offer strengths and weaknesses in terms of equipment and cost that can be tailored to the fit needs of individual labs.

## Conclusion

Through the design and optimization of the assay, it was determined that phenotypes for Growth, Resistance, and Recovery can all be accurately measured using a qPCR platform and are likely controlled by different genetic mechanisms. Existing genetic crosses can likely be used to elucidate the complex genetic interactions of the phenotypes and articulate how they may influence each other. Finally, ART-R can be effectively and accurately measured in the qPCR platform using experiment samples stored as DBS on filter paper. The methodology presented here will allow semi-high throughput implementation of the Growth, Resistance, and Recovery Assay to identify and track the emergence and spread of artemisinin partial resistance.

## List of abbreviations

ART-R: Artemisinin partial resistance
ART: Artemisinin
K13: *pfKelch13*
PC_1/2_: Patient clearance half-life
RSA: Ring-stage survival assay
DHA: Dihydroartemisinin
eRRSA: extended Recovery Ring-stage Survival Assay
GRRA: Growth, Resistance, and Recovery Assay
NaCl: Sodium Chloride
ICM: Incomplete Media
RBC: Red Blood Cell
FWB: Flow Wash Buffer
FBS: Fetal Bovine Serum
SYBR: SYBR Green I
SYTO: SYTO 61 red fluorescent nucleic acid stain
DMSO: Dimethyl sulfoxide
Ct: Cycle threshold
PMR: Parasite Multiplication Rate
DBS: Dried Blood Spot
qPCR: quantitative Polymerase Chain Reaction
ELISA: Enzyme-Linked Immunosorbent Assay

## Declarations

### Ethics approval and consent to participate

Ethical approval for the use of human blood in this study was granted by the Institutional Review Board of the University of Notre Dame. All of the blood used for the *in vitro* culturing of parasites was obtained from healthy adult volunteers and drawn by trained personnel from Interstate Blood Bank.

### Consent for publication

Not applicable.

### Availability of data and material

Data can be made available upon request to the corresponding author.

### Author’s contributions

MACS, DAS, LAC, KMB, and MTF conceived and designed the experiments. MACS, PPS, DAS, LAC, KMB, TQ, and ZC performed the experiments, analyzed the data, and created figures. MACS, DAS, LAC, JRS, and MTF wrote the manuscript. SK, XL, FHN, TCJA, and AMV contributed materials/reagents/parasites. All authors edited the manuscript. All authors read and approved the final manuscript.

## Acknowledgements

We would like to thank members of the Ferdig lab for helpful discussions.

## Additional files

**Additional file 1.**
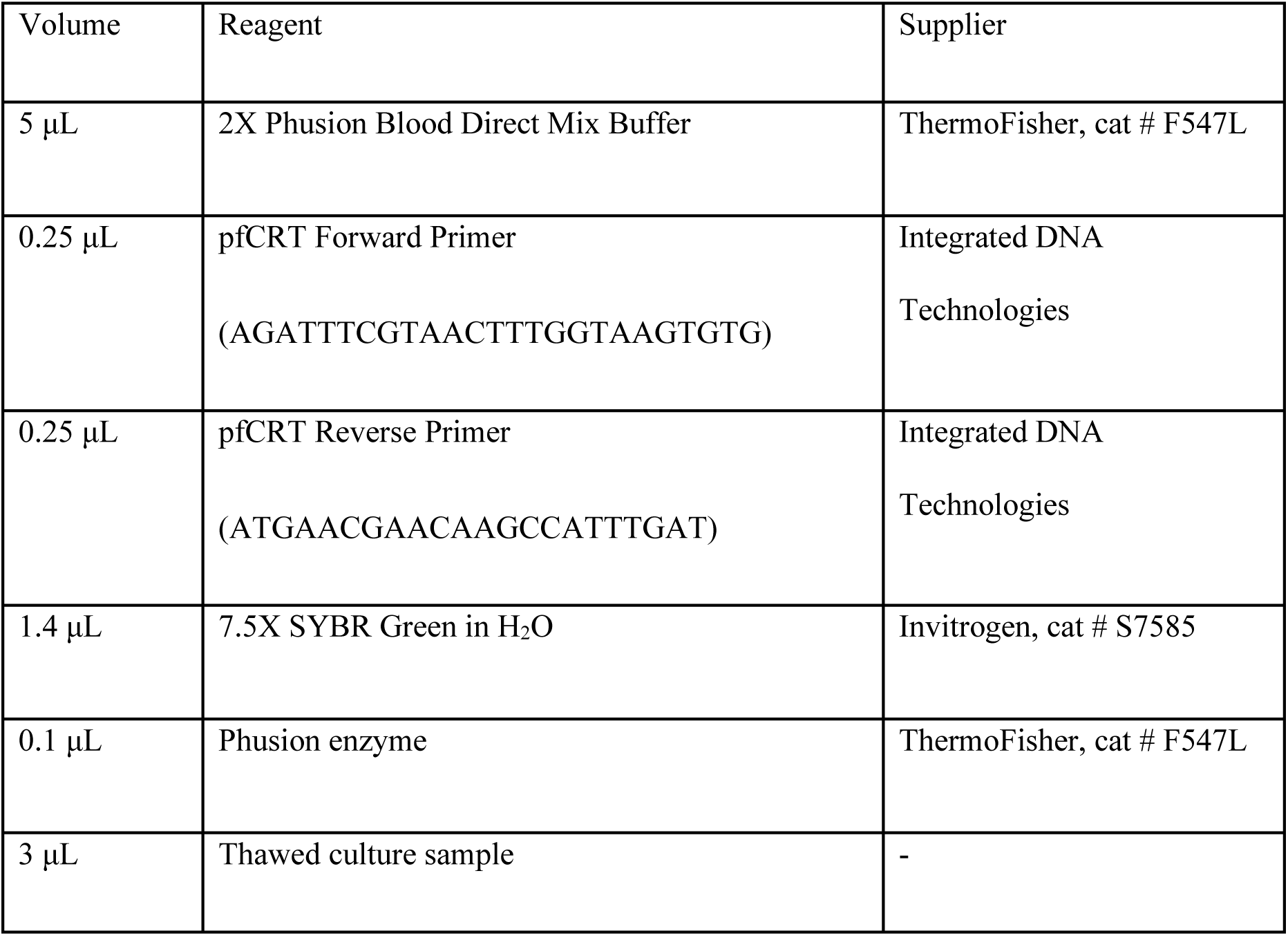
Table of primers and reagents used for quantitative PCR.

**Additional file 2.**
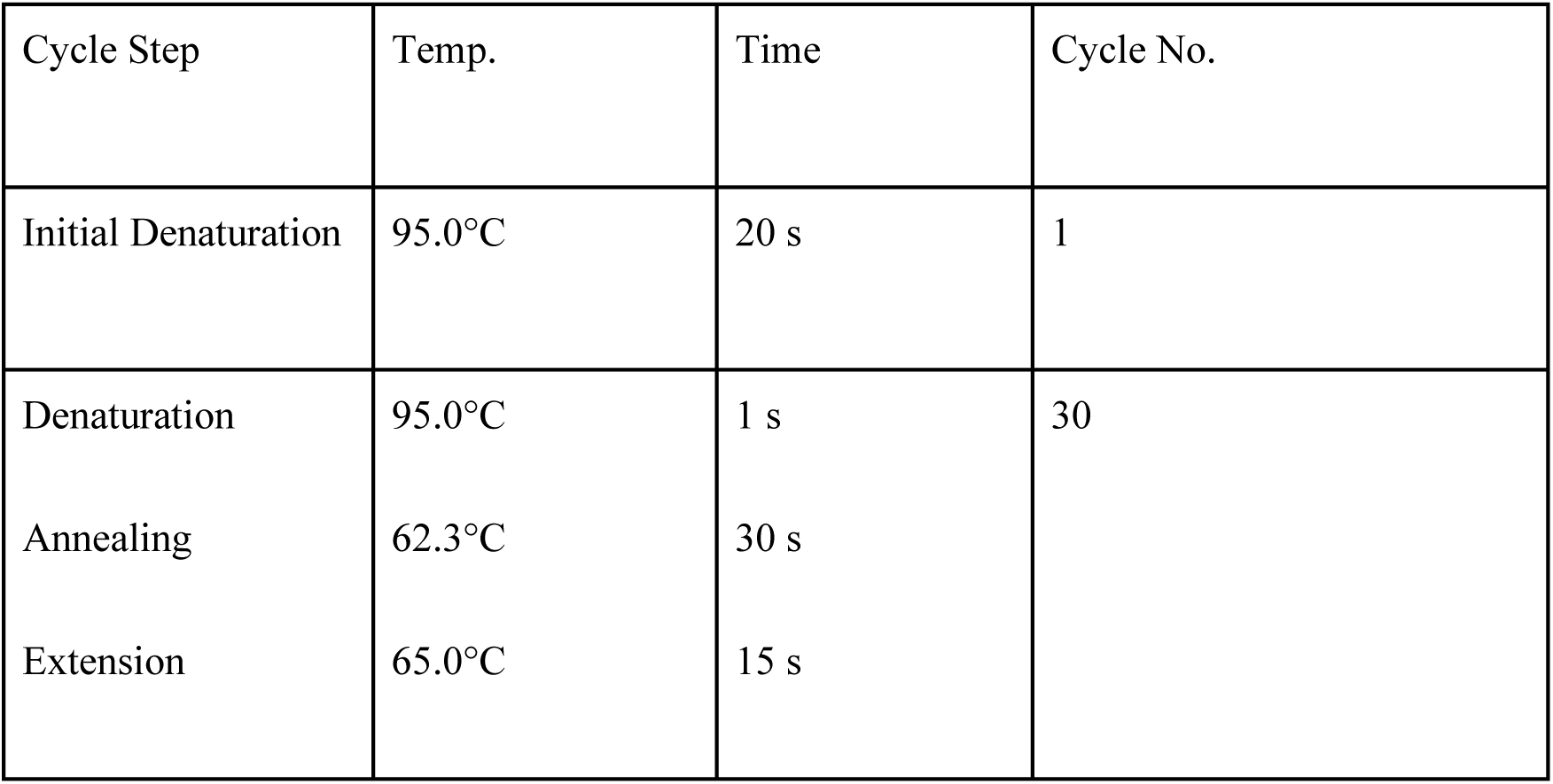
Table of cycling conditions used for quantitative PCR.

**Additional file 2.**
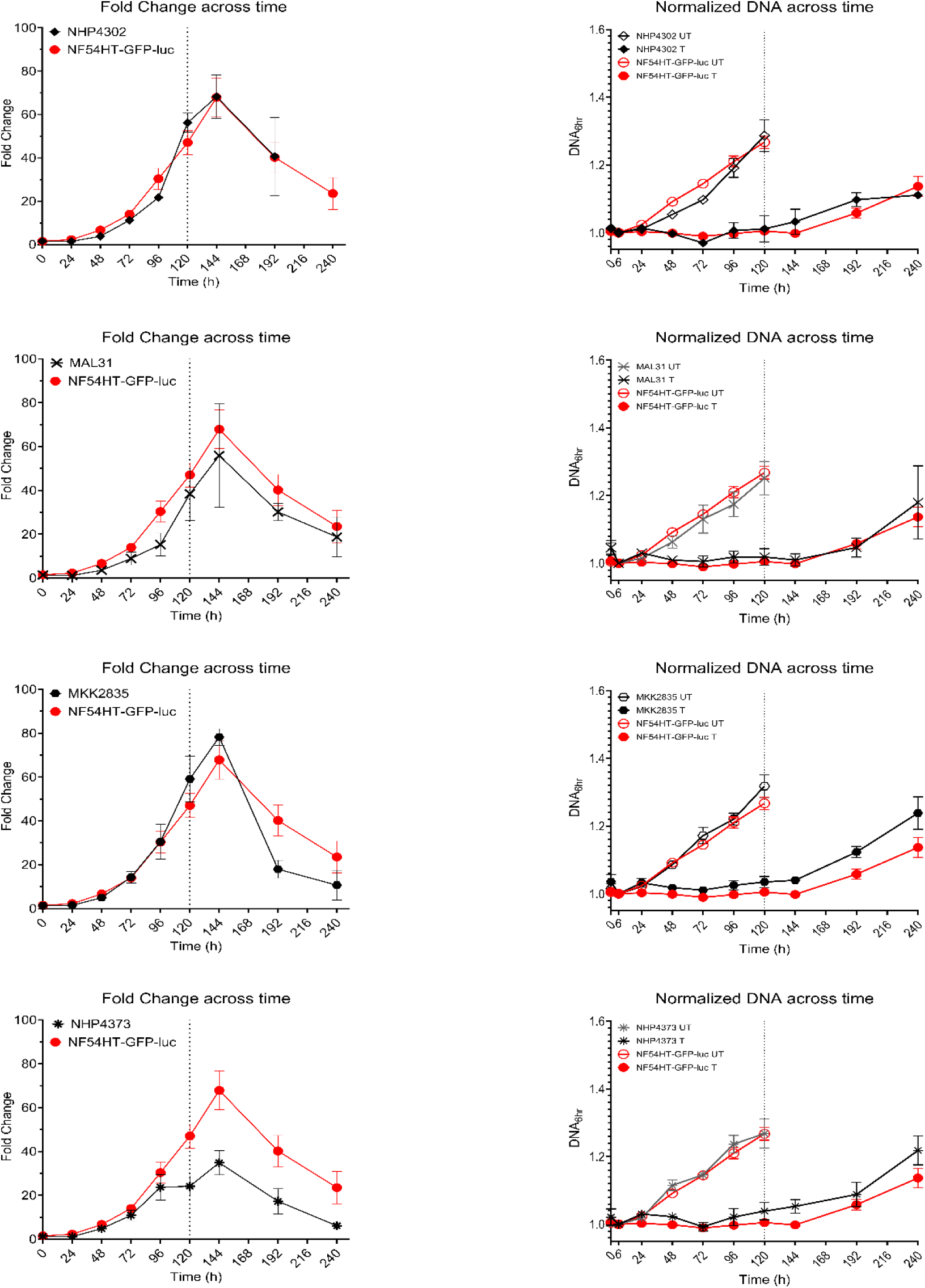
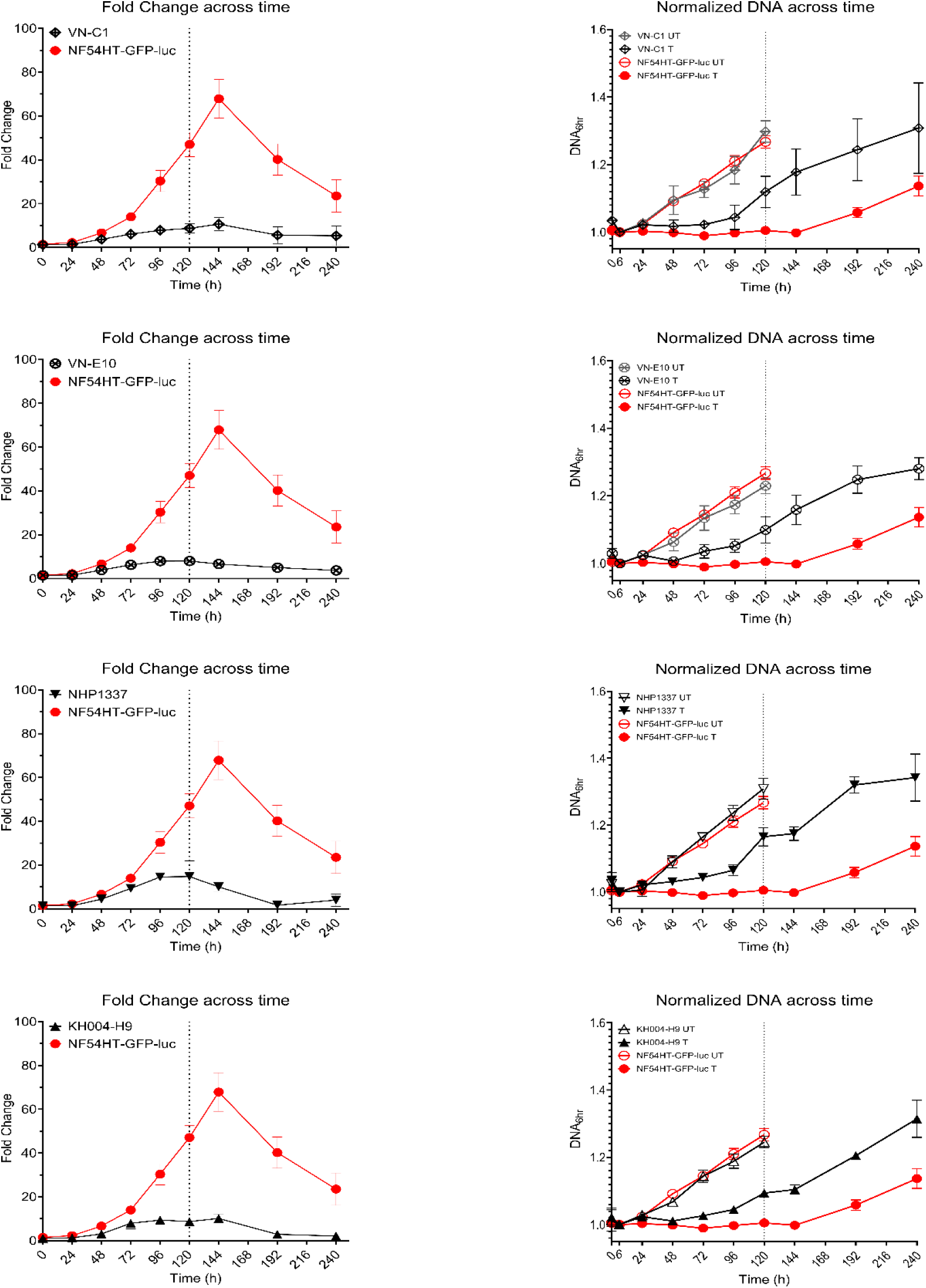
Fold Change and DNA content of treated and untreated cultures across 10-day time series for additional clinical isolates not shown in main figures. Each isolate is shown with NF54HT-GFP-luc for reference. Data is reported as mean ± SEM of at least three biological replicates.

**Additional file 3.**
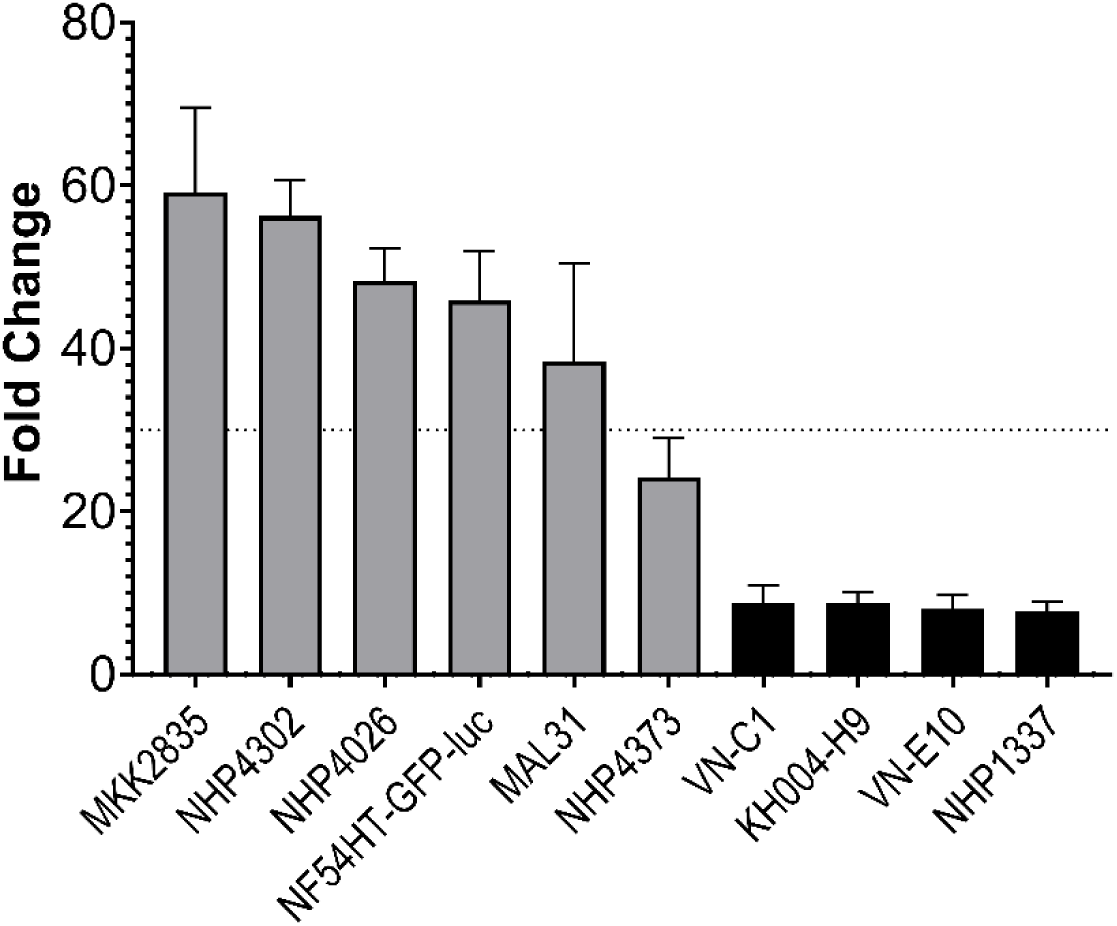
Distribution of Resistance phenotypes for clinical isolates and CRISPR/Cas9 edited K13 mutant parasite lines. Parasites are ordered by increasing level of resistance and include parents of recent genetic crosses (NF54HT-GFP-luc × NHP4026, MKK2835 × NHP1337, MAL31 × KH004-H9), NHP4302 and additional isolates (NHP4373, VN-C1, VN-E10). Parasites with a K13^C580Y^ mutation are marked in black and K13 wildtype parasites are marked in grey. The dashed line at 30-Fold Change represents the threshold for resistance, as previously reported (27). Data is reported as mean ± SEM of at least three biological replicates.

**Additional file 4.**
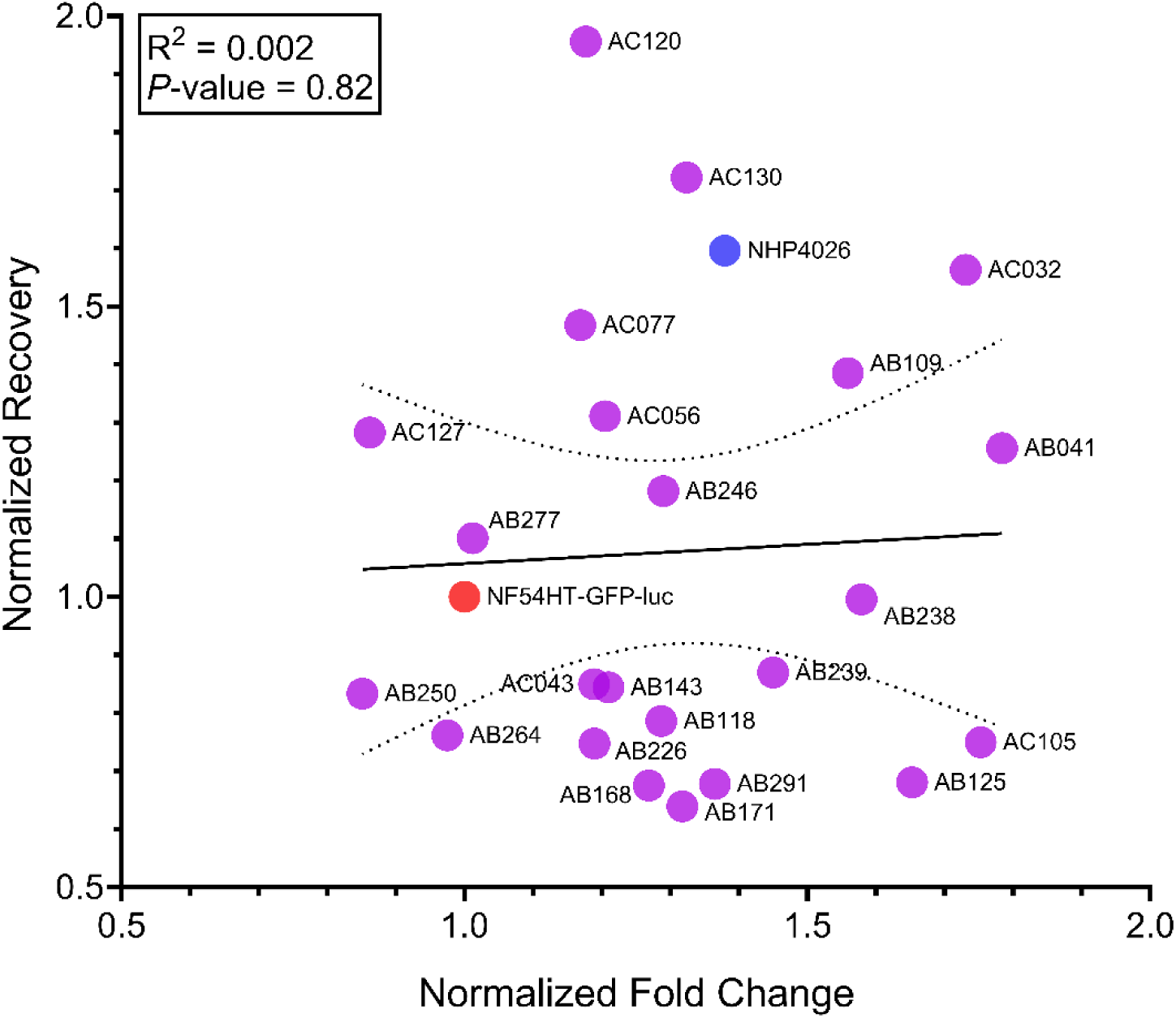
Scatterplot of Resistance vs Recovery phenotypes measured in 23 progeny and parents of the NF54(HT-GFP-luc) × NHP4026 genetic cross. Linear regression does not show a significant association Resistance and Recovery (R^2^ = 0.0021, F(1, 23) = 0.0501, *P*-value= 0.8249). The dotted lines denote the 95% confidence interval around the best fit line (*y* = 0.06616*x* + 0.9912). Linear regression was performed using mean phenotypes of at least three biological replicates and phenotype values are normalized to the NF54HT-GFP-luc parent.

**Additional file 5.**
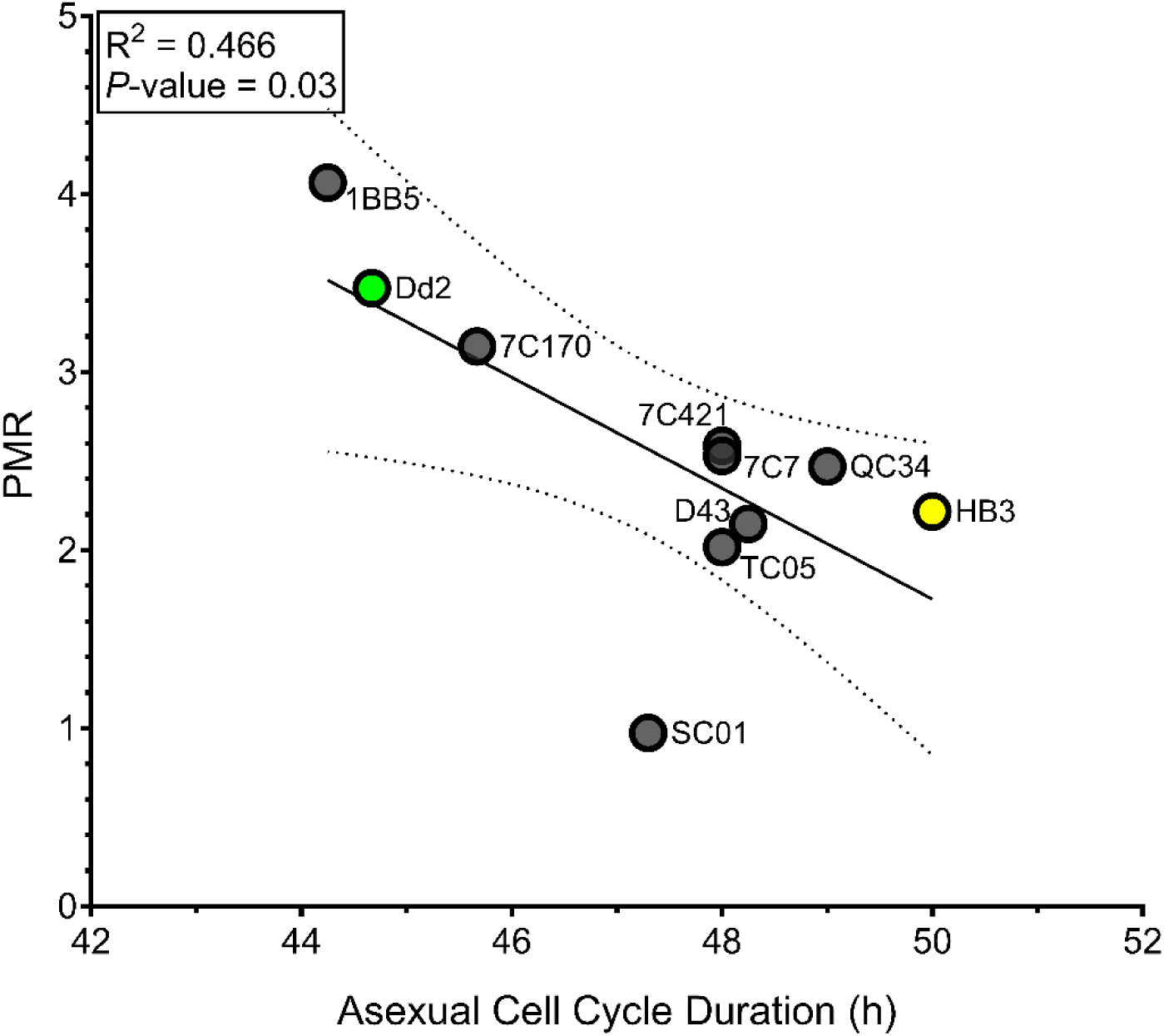
Scatterplot of PMR and asexual cell cycle duration measured in eight progeny and parents of the HB3 × Dd2 genetic cross. Linear regression between phenotypes shows a significant linear association that PMR increases as cell cycle duration decreases and 46.6% of variation is shared between the phenotypes (R^2^ = 0.4661, F(1, 8) = 6.984, *P*-value = 0.0296). The dotted lines denote the 95% confidence interval around the best fit line (*y* = - 0.3115*x* + 17.3). Linear regression was performed using mean phenotypes of at least three biological replicates or accessed from previously published sources (31).

**Additional file 6.**
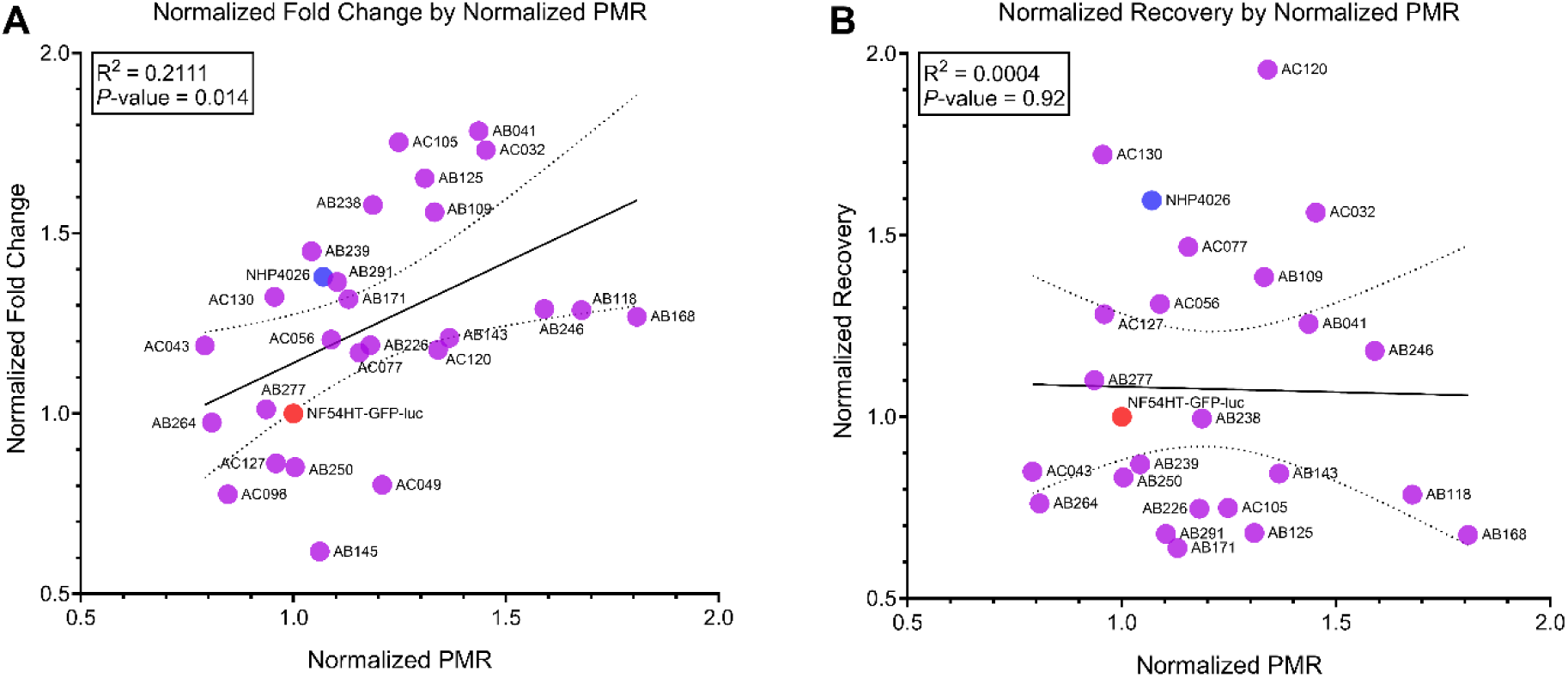
Scatterplot of PMR vs Resistance or Recovery phenotypes in progeny and parents of the NF54HT-GFP-luc × NHP4026 genetic cross. **A)** Linear regression of PMR vs Fold Change shows a significant and positive linear association with 21.1% of variation shared between the phenotypes (R^2^ = 0.2111, F(1, 26) = 6.956, *P*-value= 0.0139). The dotted lines denote the 95% confidence interval around the best fit line (*y* = 0.5582*x* + 0.5824). **B)** Linear regression of PMR vs Recovery shows no significant association (R^2^ = 0.0004, F(1, 23) = 0.009, *P*-value= 0.9234). The dotted lines denote the 95% confidence interval around the best fit line (*y* = -0.02936*x* + 1.113). Linear regression was performed using mean phenotypes of at least three biological replicates and phenotype values are normalized to the NF54HT-GFP-luc parent.

## References

1. World malaria report 2023 [Internet]. [cited 2024 Jun 23]. Available from: https://www.who.int/teams/global-malaria-programme/reports/world-malaria-report-2023

2. White NJ. Qinghaosu (Artemisinin): The Price of Success. Science. 2008 Apr 18;320(5874):330–4.

3. Ariey F, Witkowski B, Amaratunga C, Beghain J, Langlois AC, Khim N, et al. A molecular marker of artemisinin-resistant Plasmodium falciparum malaria. Nature. 2014 Jan;505(7481):50–5.

4. Takala-Harrison S, Jacob CG, Arze C, Cummings MP, Silva JC, Dondorp AM, et al. Independent Emergence of Artemisinin Resistance Mutations Among Plasmodium falciparum in Southeast Asia. The Journal of Infectious Diseases. 2015 Mar 1;211(5):670–9.

5. van Loon W, Oliveira R, Bergmann C, Habarugira F, Ndoli J, Sendegeya A, et al. In Vitro Confirmation of Artemisinin Resistance in Plasmodium falciparum from Patient Isolates, Southern Rwanda, 2019. Emerg Infect Dis. 2022 Apr;28(4):852–5.

6. Fola AA, Feleke SM, Mohammed H, Brhane BG, Hennelly CM, Assefa A, et al. Plasmodium falciparum resistant to artemisinin and diagnostics have emerged in Ethiopia. Nat Microbiol. 2023 Oct;8(10):1911–9.

7. Conrad MD, Asua V, Garg S, Giesbrecht D, Niaré K, Smith S, et al. Evolution of Partial Resistance to Artemisinins in Malaria Parasites in Uganda. New England Journal of Medicine. 2023 Aug 24;389(8):722–32.

8. Dondorp AM, Nosten F, Yi P, Das D, Phyo AP, Tarning J, et al. Artemisinin resistance in Plasmodium falciparum malaria. N Engl J Med. 2009 Jul 30;361(5):455–67.

9. Flegg JA, Guerin PJ, White NJ, Stepniewska K. Standardizing the measurement of parasite clearance in falciparum malaria: the parasite clearance estimator. Malaria Journal. 2011;10(1):339.

10. Vestergaard LS, Ringwald P. Responding to the challenge of antimalarial drug resistance by routine monitoring to update national malaria treatment policies. Am J Trop Med Hyg. 2007 Dec;77(6 Suppl):153–9.

11. Flegg JA, Guérin PJ, Nosten F, Ashley EA, Phyo AP, Dondorp AM, et al. Optimal sampling designs for estimation of Plasmodium falciparum clearance rates in patients treated with artemisinin derivatives. Malar J. 2013 Nov 13;12(1):411.

12. Ataide R, Ashley EA, Powell R, Chan JA, Malloy MJ, O’Flaherty K, et al. Host immunity to Plasmodium falciparum and the assessment of emerging artemisinin resistance in a multinational cohort. Proceedings of the National Academy of Sciences. 2017 Mar 28;114(13):3515–20.

13. Chotivanich K, Udomsangpetch R, McGready R, Proux S, Newton P, Pukrittayakamee S, et al. Central Role of the Spleen in Malaria Parasite Clearance. The Journal of Infectious Diseases. 2002 May 15;185(10):1538–41.

14. Witkowski B, Amaratunga C, Khim N, Sreng S, Chim P, Kim S, et al. Novel phenotypic assays for the detection of artemisinin-resistant Plasmodium falciparum malaria in Cambodia: in-vitro and ex-vivo drug-response studies. The Lancet Infectious Diseases. 2013 Dec 1;13(12):1043–9.

15. Mukherjee A, Bopp S, Magistrado P, Wong W, Daniels R, Demas A, et al. Artemisinin resistance without pfkelch13 mutations in Plasmodium falciparum isolates from Cambodia. Malaria Journal. 2017 May 12;16(1):195.

16. Balikagala B, Fukuda N, Ikeda M, Katuro OT, Tachibana SI, Yamauchi M, et al. Evidence of Artemisinin-Resistant Malaria in Africa. New England Journal of Medicine. 2021 Sep 23;385(13):1163–71.

17. Noedl Harald, Se Youry, Schaecher Kurt, Smith Bryan L., Socheat Duong, Fukuda Mark M. Evidence of Artemisinin-Resistant Malaria in Western Cambodia. New England Journal of Medicine. 2008;359(24):2619–20.

18. Amaratunga C, Sreng S, Suon S, Phelps ES, Stepniewska K, Lim P, et al. Artemisinin-resistant Plasmodium falciparum in Pursat province, western Cambodia: a parasite clearance rate study. The Lancet Infectious Diseases. 2012 Nov;12(11):851–8.

19. Chotivanich K, Tripura R, Das D, Yi P, Day NPJ, Pukrittayakamee S, et al. Laboratory Detection of Artemisinin-Resistant Plasmodium falciparum. Antimicrobial Agents and Chemotherapy. 2014 May 14;58(6):3157–61.

20. Witkowski B, Khim N, Chim P, Kim S, Ke S, Kloeung N, et al. Reduced Artemisinin Susceptibility of Plasmodium falciparum Ring Stages in Western Cambodia. Antimicrobial Agents and Chemotherapy. 2013 Jan 22;57(2):914–23.

21. Witkowski B, Lelièvre J, Barragán MJL, Laurent V, Su X zhuan, Berry A, et al. Increased Tolerance to Artemisinin in Plasmodium falciparum Is Mediated by a Quiescence Mechanism. Antimicrobial Agents and Chemotherapy. 2010 May 1;54(5):1872–7.

22. Reyser T, Paloque L, Ouji M, Nguyen M, Ménard S, Witkowski B, et al. Identification of compounds active against quiescent artemisinin-resistant Plasmodium falciparum parasites via the quiescent-stage survival assay (QSA). Journal of Antimicrobial Chemotherapy. 2020 Oct 1;75(10):2826–34.

23. Teuscher F, Gatton ML, Chen N, Peters J, Kyle DE, Cheng Q. Artemisinin-Induced Dormancy in Plasmodium falciparum: Duration, Recovery Rates, and Implications in Treatment Failure. The Journal of Infectious Diseases. 2010 Nov 1;202(9):1362–8.

24. Hott A, Casandra D, Sparks KN, Morton LC, Castanares GG, Rutter A, et al. Artemisinin-Resistant Plasmodium falciparum Parasites Exhibit Altered Patterns of Development in Infected Erythrocytes. Antimicrobial Agents and Chemotherapy. 2015 Jun;59(6):3156–67.

25. Watts RE, Odedra A, Marquart L, Webb L, Abd-Rahman AN, Cascales L, et al. Safety and parasite clearance of artemisinin-resistant Plasmodium falciparum infection: A pilot and a randomised volunteer infection study in Australia. PLOS Medicine. 2020 Aug 21;17(8):e1003203.

26. Peatey C, Chen N, Gresty K, Anderson K, Pickering P, Watts R, et al. Dormant Plasmodium falciparum Parasites in Human Infections Following Artesunate Therapy. The Journal of Infectious Diseases. 2021 May 1;223(9):1631–8.

27. Davis SZ, Singh PP, Vendrely KM, Shoue DA, Checkley LA, McDew-White M, et al. The extended recovery ring-stage survival assay provides a superior association with patient clearance half-life and increases throughput. Malaria Journal. 2020 Jan 31;19(1):54.

28. Vendrely KM, Kumar S, Li X, Vaughan AM. Humanized Mice and the Rebirth of Malaria Genetic Crosses. Trends in Parasitology. 2020 Oct 1;36(10):850–63.

29. Vaughan AM, Pinapati RS, Cheeseman IH, Camargo N, Fishbaugher M, Checkley LA, et al. Plasmodium falciparum genetic crosses in a humanized mouse model. Nat Methods. 2015 Jul;12(7):631–3.

30. Amambua-Ngwa A, Button-Simons KA, Li X, Kumar S, Brenneman KV, Ferrari M, et al. Chloroquine resistance evolution in Plasmodium falciparum is mediated by the putative amino acid transporter AAT1. Nat Microbiol. 2023 Jul;8(7):1213–26.

31. Reilly Ayala HB, Wacker MA, Siwo G, Ferdig MT. Quantitative trait loci mapping reveals candidate pathways regulating cell cycle duration in Plasmodium falciparum. BMC Genomics. 2010;11(1):577.

32. Dluzewski AR, Ling IT, Rangachari K, Bates PA, Wilson RJM. A simple method for isolating viable mature parasites of Plasmodium falciparum from cultures. Transactions of the Royal Society of Tropical Medicine and Hygiene. 1984 Jan 1;78(5):622–4.

33. Mancio-Silva L, Lopez-Rubio JJ, Claes A, Scherf A. Sir2a regulates rDNA transcription and multiplication rate in the human malaria parasite Plasmodium falciparum. Nat Commun. 2013 Feb 26;4(1):1530.

34. Taioli E, Kinney P, Zhitkovich A, Fulton H, Voitkun V, Cosma G, et al. Application of reliability models to studies of biomarker validation. Environmental Health Perspectives. 1994 Mar;102(3):306–9.

35. Tirrell AR, Vendrely KM, Checkley LA, Davis SZ, McDew-White M, Cheeseman IH, et al. Pairwise growth competitions identify relative fitness relationships among artemisinin resistant Plasmodium falciparum field isolates. Malar J. 2019 Aug 28;18(1):295.

36. Reilly HB, Wang H, Steuter JA, Marx AM, Ferdig MT. Quantitative dissection of clone-specific growth rates in cultured malaria parasites. International Journal for Parasitology. 2007 Dec 1;37(14):1599–607.

37. Button-Simons KA, Kumar S, Carmago N, Haile MT, Jett C, Checkley LA, et al. The power and promise of genetic mapping from Plasmodium falciparum crosses utilizing human liver-chimeric mice. Commun Biol. 2021 Jun 14;4(1):1–13.

38. Phyo AP, Nkhoma S, Stepniewska K, Ashley EA, Nair S, McGready R, et al. Emergence of artemisinin-resistant malaria on the western border of Thailand: a longitudinal study. The Lancet. 2012 May;379(9830):1960–6.

39. Teuscher F, Chen N, Kyle DE, Gatton ML, Cheng Q. Phenotypic Changes in Artemisinin-Resistant Plasmodium falciparum Lines In Vitro: Evidence for Decreased Sensitivity to Dormancy and Growth Inhibition. Antimicrobial Agents and Chemotherapy. 2012 Dec 26;56(1):428–31.

40. Fukuda N, Yoshida N, Balikagala B, Tsuru I, Ikeda M, Hirai M, et al. Detection of drug-resistant malaria in resource-limited settings: efficient and high-throughput surveillance of artemisinin and partner drug resistance. Journal of Antimicrobial Chemotherapy. 2024 Apr 25;dkae120.

41. Brenneman KV, Li X, Kumar S, Delgado E, Checkley LA, Shoue DA, et al. Optimizing bulk segregant analysis of drug resistance using Plasmodium falciparum genetic crosses conducted in humanized mice. iScience. 2022 Apr 15;25(4):104095.

42. Kane J, Li X, Kumar S, Button-Simons KA, Vendrely Brenneman KM, Dahlhoff H, et al. A Plasmodium falciparum genetic cross reveals the contributions of pfcrt and plasmepsin II/III to piperaquine drug resistance. mBio. 2024 Jun 24;0(0):e00805–24.

